# Secretome of senescent hepatoma cells modulate macrophage polarization and neutrophil extracellular traps formation

**DOI:** 10.1101/2020.12.30.423905

**Authors:** Bijoya Sen, Savera Aggarwal, Rhisita Nath, Rashi Sehgal, Archana Rastogi, Nirupma Trehanpati, Gayatri Ramakrishna

## Abstract

Presence of dysfunctional senescent hepatocyte is a hallmark feature of cirrhosis. We now report the presence of senescent hepatocytes (p21 and p53 positive) in vicinity of infiltrated immune cells in hepatocellular carcinoma. Hence, we checked if senescent cells can alter fate of macrophage polarization and neutrophil extracellular trap (NETs) formation. Using an in vitro assay, senescence was induced in hepatoma cells (HepG2 and Huh7 cells) by doxorubicin treatment and senescent cell showed secretory phenotype with strong expression of cytokines (IL1β, IL6, IL8 and IL13) as evaluated by Flow cytometry. The senescent secretome from hepatoma cells induced macrophage differentiation predominantly with M2 markers (CD80, CD86) while that of non-senescent cell induced M1 phenotype (CD163, CD206) as analysed by flow cytometry. Human hepatocellular carcinoma harbouring senescent hepatocytes showed presence of M2 macrophages, while M1 macrophages were predominant in non-tumorous region. Additionally, the senescent secretome from Huh7 cells (p53^mut^) enhanced the NETs formation, while HepG2 (p53^+/+^) secretome had an inhibitory affect In conclusion, the “pro-inflammatory” senescent secretome drives non-inflammatory type M2 macrophage polarization and modulate neutrophil traps thereby modulating the microenvironment towards tumor promotion. Targeting senescent hepatocyte secretome appears a promising therapeutic target in liver cancer in future.

Work Highlight (Diagrammatic Representation).

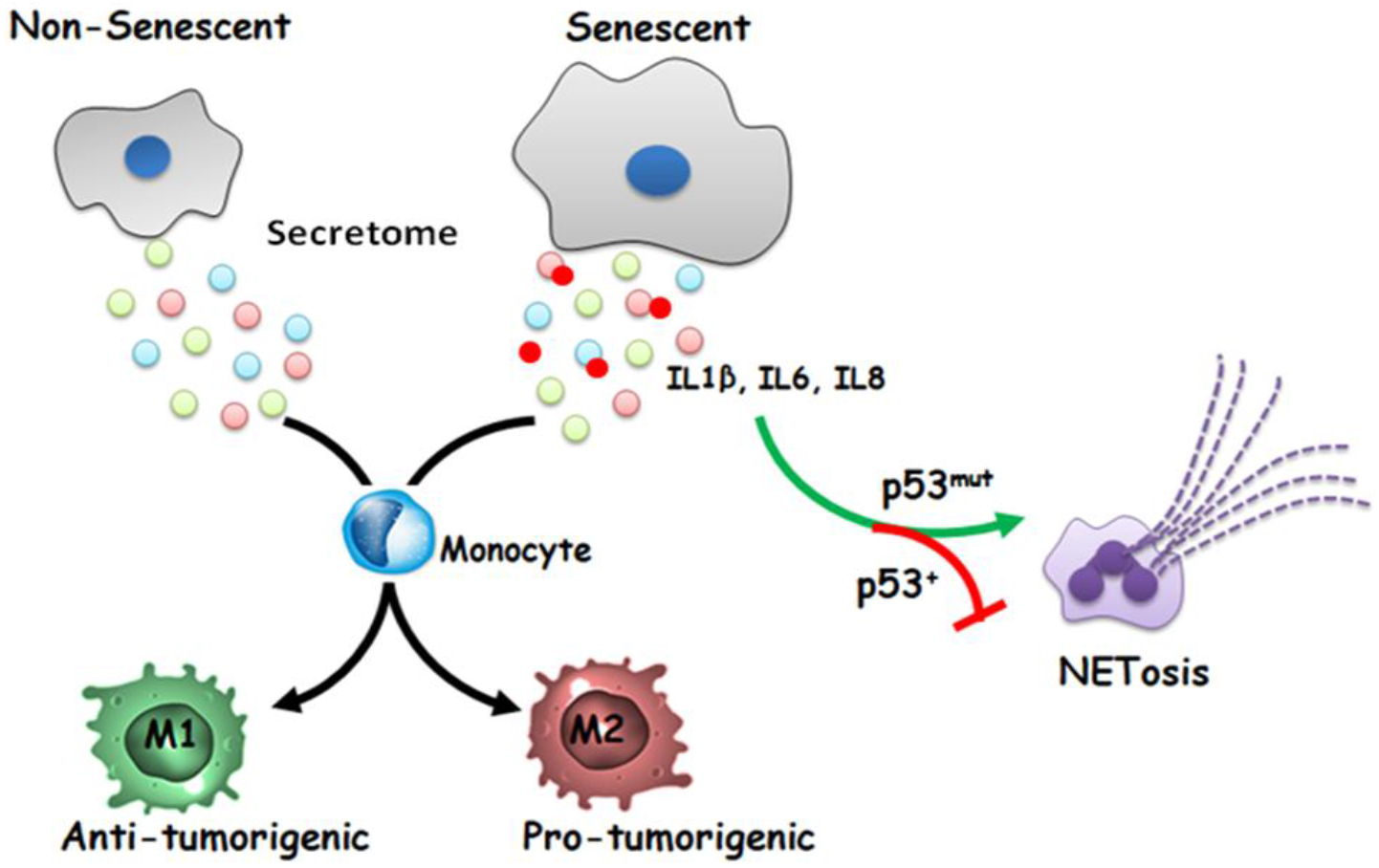

## Introduction

Cellular senescence is a state of permanent growth arrest and serves as a double sword edge in progression of cancer. Senescence acts as barrier for tumorigenesis as senescent cells do not proliferate, while a pro-inflammatory senescence-associated secretory phenotype (SASP) may provide a conducive milieu to drive neoplasia. Hepatocellular carcinoma usually originates on a cirrhotic background which is bed of chronic inflammation [1–4]. Senescent hepatocytes cells have been often noted in cirrhosis [5–8]. We recently showed accumulation of senescent hepatocytes in decompensated cirrhosis and further it is associated with downregulation of mitochondrial unfolded protein response [9].

Chemotherapeutic agents can induce senescence even in the cancer cells and often the senescent cells persist for a long time [10]. A characteristic feature of senescent cells is its senescence associated secretory phenotype (SASP) which can help in chemoattraction of immune cells and remodel the extracellular matrix. However, the impact of senescent cells on the modulation of innate immune cell fate in tumor microenvironment is less explored. Senescent hepatocytes can promote migration of macrophages [11] and during liver fibrosis, the senescent stellate cells can influence the macrophages [12].

The hepatic infiltration of neutrophils is essential for an active defence against the invading microorganisms. However, an overt activation of neutrophil can cause tissue damage by the phenomenon of neutrophil extracellular traps (NETs) whereby neutrophils release their decondensed chromatin and granules into extracellular space. Recently role of NETs has been described in liver steatosis and inhibiting NETs formation is linked with decreased tumor growth [13]. There is no information available of role of senescent secretome on NETs formation.

As hepatocyte senescence is a hallmark features of cirrhosis [8,9,14], and often hepatocellular carcinoma occurs on background of cirrhosis, we hypothesized that senescent cell can influence the fate of HCC by its secretome. It has been reported that cytokines secreted by senescent cells can directly promote proliferation of susceptible cancer cells via paracrine signalling [15–17]. However, the information is scant with respect to existence of senescent hepatocyte in liver tumor and how its secretome can influence the infiltrating immune cells. Therefore, in the present study, we have explored the whether secretome of senescent cells effect the innate immune cell fate viz., macrophage polarization and neutrophil extracellular trap formation.

## Materials and Methods

### Induction of premature senescence and collection of senescent secretome

HepG2 and Huh7 cells were purchased cell repository at National Cell Science Centre, Pune (India) and were all mycoplasma negative. Senescence was induced by treating with doxorubicin as described by us [9] using 2μM Doxorubicin (Dox) (Sigma, St. Louis, MO) and senescence was confirmed by Senescence associated-β-Galactosidase assay [18]. The expression of the inflammatory markers were checked by Flow cytometry. For collection of secretome, on the fifth day of treatment, media was changed with 70:30 ratio of RPMI:DMEM supplemented with 10% FBS and senescent secretome was collected fter 24 hrs. The collected secretome was clarified by high speed centrifugation, aliquot and kept in −80°C until use.

### Human tissue samples

Stored formalin fixed paraffin embedded tissue samples were obtained from the pathology and the blood samples from healthy subjects were obtained after informed consent. The study was approved from ILBS institutional ethics committee (approval # F25/5/43/AC/ILBS/2013/1157) according to Helinski declaration.

### Immunohistochemistry

This was performed on formalin fixed paraffin embedded tissue sections (4μM) as described earlier [9]. Briefly the tissue was deparaffinised, followed by antigen retrieval, power block and antibody additions. The later steps were performed as described by the BioGenex Kit (Fremont, CA). The stained sections were read by pathologists.

### Immunofluorescence

Cells plated on coverslips were fixed with 4% formaldehyde, blocked in 3% BSA followed by incubation with primary antibody overnight at 4°C. For tissue samples, 4 μM FFPE tissue sections were antigen retrieved and incubated with primary antibody overnight at 4°C. Primary antibody was detected using Alexa Fluor 488 or 594 conjugated secondary (anti-mouse/rabbit/goat) antibodies and mounted in DAPI mounting media.

### Isolation of peripheral blood mononuclear cells (PBMCs) and polymorphonuclear granulocytes (PMNs)

Blood was collected from healthy donors (N=5) and PBMCs were isolated using Lymphoprep™ (Alere Technologies AS, Oslo, Norway) according to manufacturer’s protocol. PMNs were isolated using Polymorphprep™ (Alere Technologies AS, Oslo, Norway) according to manufacturer’s protocol. About 80% of polymorphonuclear granulocytes mostly contained neutrophils as checked by Lesihman stain.

### Induction of macrophage polarisation and their flow cytometry analysis

Isolated PBMCs from a single individual were divided into parts and cultured in presence of either control secretome, senescent secretome, control media or with positive controls: GM-CSF (10ng/ml) for M1 or MCSF (10ng/ml) for M2. Cells were cultured for 6 days with media change on every alternate day. On 6^th^ day, cells were trypsinised and recovered for surface markers in media containing 10% FBS for 3h in CO_2_ incubator at 37°C. Cell surface markers were identified with CD68-FITC (eBioscience, San Diego, CA), CD80-V450 and CD86-PEcy7 (BD Biosciences, San Jose, CA) as M1 markers and CD206-PEcy5 and CD163-APC (BD Biosciences, San Jose, CA) as M2 markers by using flow cytometry. The baseline parameters of the same individual were also checked on 0 day of treatment i.e. on the day of PBMC isolation and used for comparison. The frequencies of the different subpopulations were calculated using FlowJo^®^ software (Beckton-Dickenson, Franklin Lakes, NJ).

### NETosis induction and estimation

Isolated PMNs (5 × 10^5^) which consisted majority of the neutrophils were treated with either 500 μl of control secretome, senescent secretome, control media or with PMA (50ng/ml). Impermeable DNA dye, Sytox Green (λ_em_: 523nm, Invitrogen, Carlsbad, CA) was added to each tube to a final concentration of 0.2 μM. Cell permeable Hoechst 33258 (λ_em_: 461nm, Cayman Chemical, Ann Arbor, MI) was also added at working concentration of 2μg/ml. 100μl of stained cells were transferred to 96-well black plate in quadruplets and then incubated for 4h in CO_2_ incubator. Fluorescence was quantified in FLUOstar OPTMA (BMG LABTECH, Ortenberg, Germany) by using λ_ex/em_ of 355/460 for Hoechst and 485/520 for Sytox Green. Results were calculated as relative fluorescence units (RFU = SYTOX Green/Hoechst × 100) and were expressed as relative fold change by dividing the average of the experimental condition (secretome) by the average of the untreated condition (RPMI:DMEM) [19].

## Results

### Presence of senescent hepatocytes in hepatocellular carcinoma

Previously we had reported accumulation of senescent cells in cirrhosis [9], however the presence of senescent cells in HCC was not known. The cirrhotic liver showed intense positivity for senescence markers p21 and p53 and feeble expression of proliferation marker Ki67 (shown in Fig. 1A). As expected the HCC specimens showed high proliferative index as seen by Ki67 staining and the positivity for p21 and p53 were extremely low in both the normal and HCC specimens (shown in Fig. 1A, B). Inspite of the low number of senescent cells (p21/p53 positive) in HCC, an intriguing observation was that the senescent hepatocytes (p21 or p53 positive cells) existed in close vicinity of immune cells (Fig.1A). However, same was not seen in the normal liver sections. Hence, in the subsequent work we evaluated if senescent hepatocytes can affect the fate of the immune cells.

**Fig. 1.**
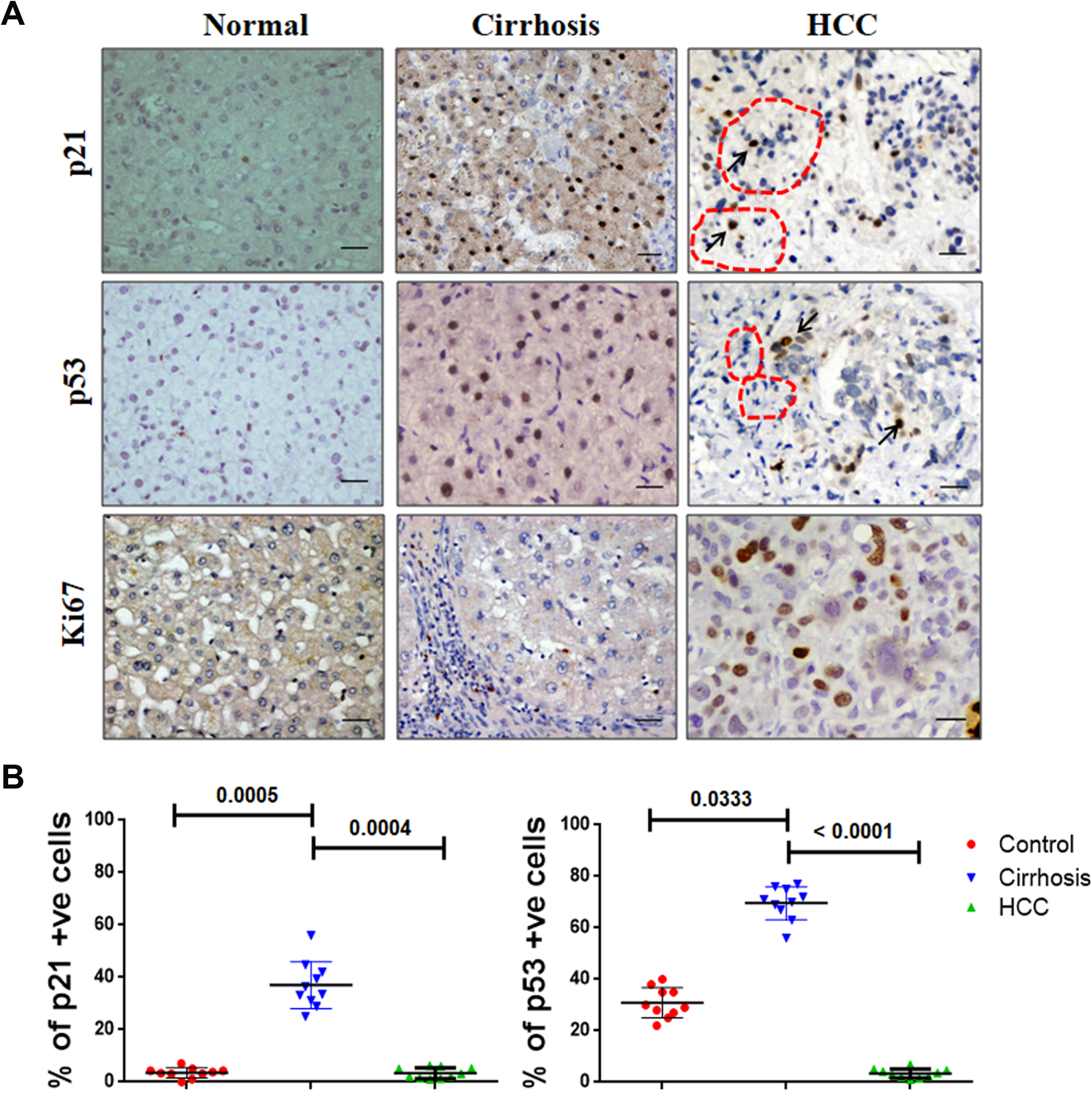
Immunohistochemistry for senescence associated growth arrested markers viz., p53 and p21, together with the proliferative marker, Ki67 in various stages of liver disease. (A) Brown staining indicates positivity. Red dotted lines represent infiltrating immune cells adjacent to senescent hepatocytes. Black arrows indicate p21 and p53 positive senescent hepatocytes in HCC. (B) Scatter plots showing changes in percentage positivity of p21 and p53. Occasional (1-5%) senescent hepatocytes were also noted in HCC. Data represents mean ± SEM. Kruskal Wallis One way ANOVA was used to determine the statistical significance.

### Doxorubicin induced premature senescence is associated with an inflammatory phenotype

Senescence was induced in hepatocellular carcinoma cells (Huh7 and HepG2 cells) by treating with low dose of doxorubicin as described by us earlier [9]. In brief, low dose of doxorubicin induced typical features of senescence such as SA-β galactosidase positivity, loss of Lamin B1 and growth inhibition in S-phase as indicated by EdU labelling (shown in Fig. S1A-D). In the subsequent work we have now referred the doxorubicin treated cells as “senescent cells”. Next the hallmark feature of senescence-associated secretory phenotype was checked by flow cytometry. The level of the cytokines, IL1β, IL6, IL8 and IL13 were significantly increased in senescent hepatoma cells compared to the control cells (shown in Fig. 2A). Interestingly, Huh7 cells with a mutant p53 showed higher basal level of the cytokines compared to HepG2 cells with wild type p53 status and corroborates finding on increased p53 activity reduces inflammatory cytokine production [20]. Premature senescence in hepatoma cell lines was also accompanied with an increase in frequency of cells expressing CCR7 and CCL4 which have been previously shown to cause chemoattraction of various immune cells such as macrophages, NK cells, and T cells [21] (shown in Fig. 2B).

**Fig. 2.**
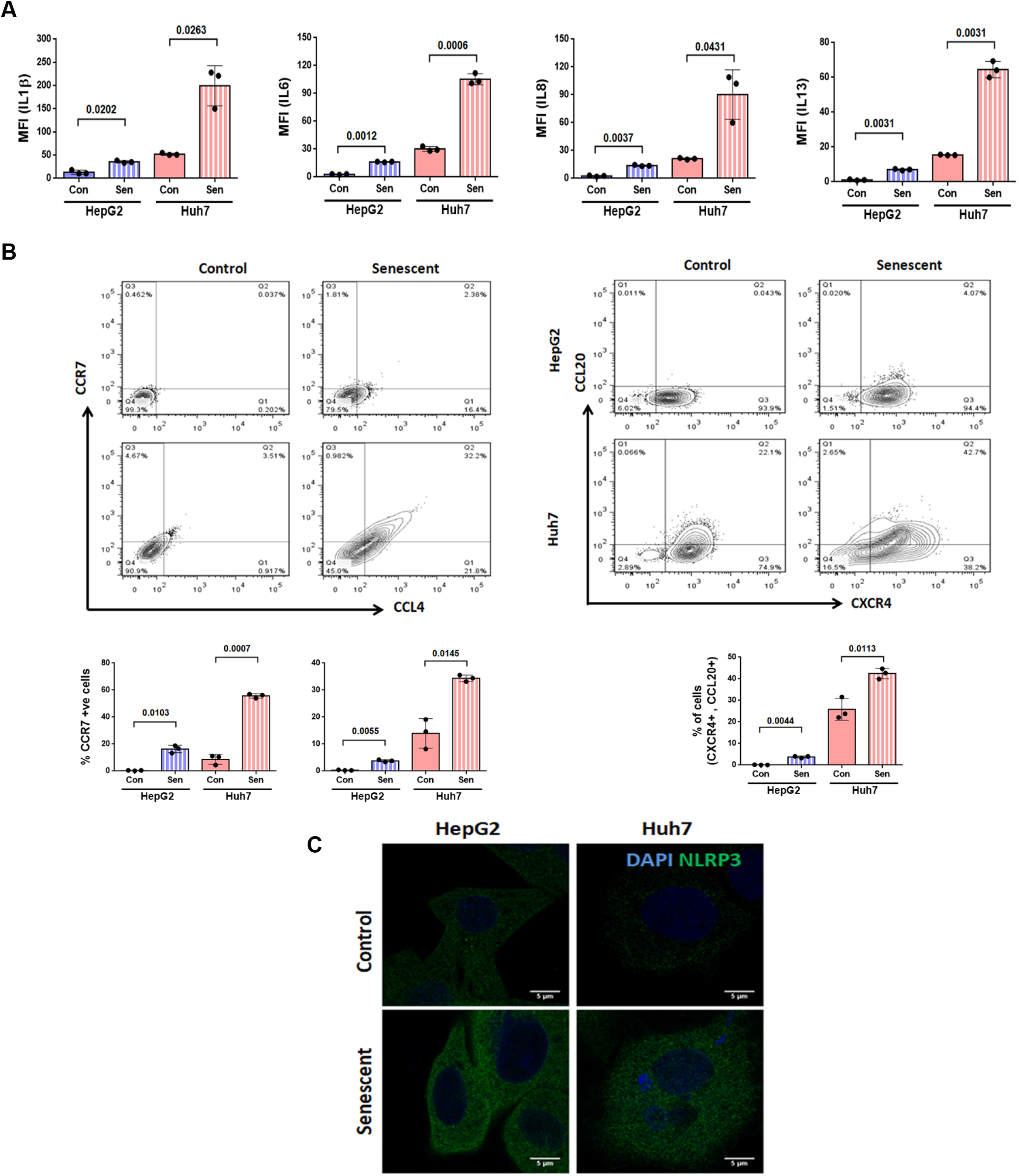
Senescent hepatoma cells are inflammatory in nature. (A) Flow cytometry analysis of inflammatory cytokines, IL1β, IL6, IL8 and IL13 in control (Con) and senescent (Sen) HepG2 and Huh7 cells. The bar graph shows median fluorescent intensity (MFI) of cytokines. (B) Flow cytometry analysis for the expression of CCR7, CCL4, CCL20 and CXCR4 in control and senescent hepatoma cells. The top panel shows representative contour plots and the bottom panel shows the percentage positive cells for representative markers. Individual data points represent biological replicates and data is represented as mean ± SD. P values were calculated by Student’s t test. (C) Merged confocal images showing expression of NLRP3 (green) in control and senescent HepG2 and Huh7 cells. Cells were counterstained with DAPI (blue).

Additionally, higher frequency of cells with CCL20 and CXCR4 expression was noted in senescent cells (shown in Fig. 2B) suggestive of its possible influence on increased tumor growth and invasiveness [22]. Since inflammasome complex play an important role in the maturation of proinflammatory cytokine IL1β, The expression level of NLRP3 was increased in senescent HepG2 and Huh7 cells (Fig. 2C) indicating its role of inflammasome [23]. The above results indicated that pro-inflammatory SASP is a key characteristic feature of senescent hepatoma cells.

### Senescent secretome drives M2 type macrophage polarisation

As senescent cells are in vicinity of immune cells in hepatocellular carcinoma, we asked the question whether the senescent secretome influences the immune cell fate. PBMCs were cultured in presence of senescent secrteome and analyzed for macrophage polarization. The cultured monocytes/macrophages were identified according to side scatter and CD68 profile and subsequently expression of CD80, CD86, CD163 and CD206 were analyzed to delineate the M1 and M2 macrophages (3 A). Briefly, M1 were classified as cells with high expression of CD80, CD86, while macrophages showing high expression of CD163, CD206 were considered as M2 subtype (Fig. 3B). The results indicated that the senescent secretome drives the macrophages towards protumorigenic M2 type with increase in frequency of cells expressing CD206 and CD163 compared to control secretome which induced polarization towards the M1 type with increased frequency of cells expressing CD80 and CD86 (shown in Fig. 3C). Macrophage polarization was also checked in HCC, by lineage specific markers for M1 and M2 i.e. iNOS and arginase1 respectively in tumor and non-tumor areas. The tumor area showed CD68 positive macrophages with arginase1 expression indicative of predominance of M2 macrophages; whereas, the macrophages in non-tumor area showed iNOS positivity indicative of M1 polarized cells (shown in Fig. 3D). Overall, the above *in vitro* results indicate that SASP from senescent hepatoma cells drive alternatively activated M2 type macrophage polarisation which may possibly promote tumor growth and invasiveness [24].

**Fig. 3.**
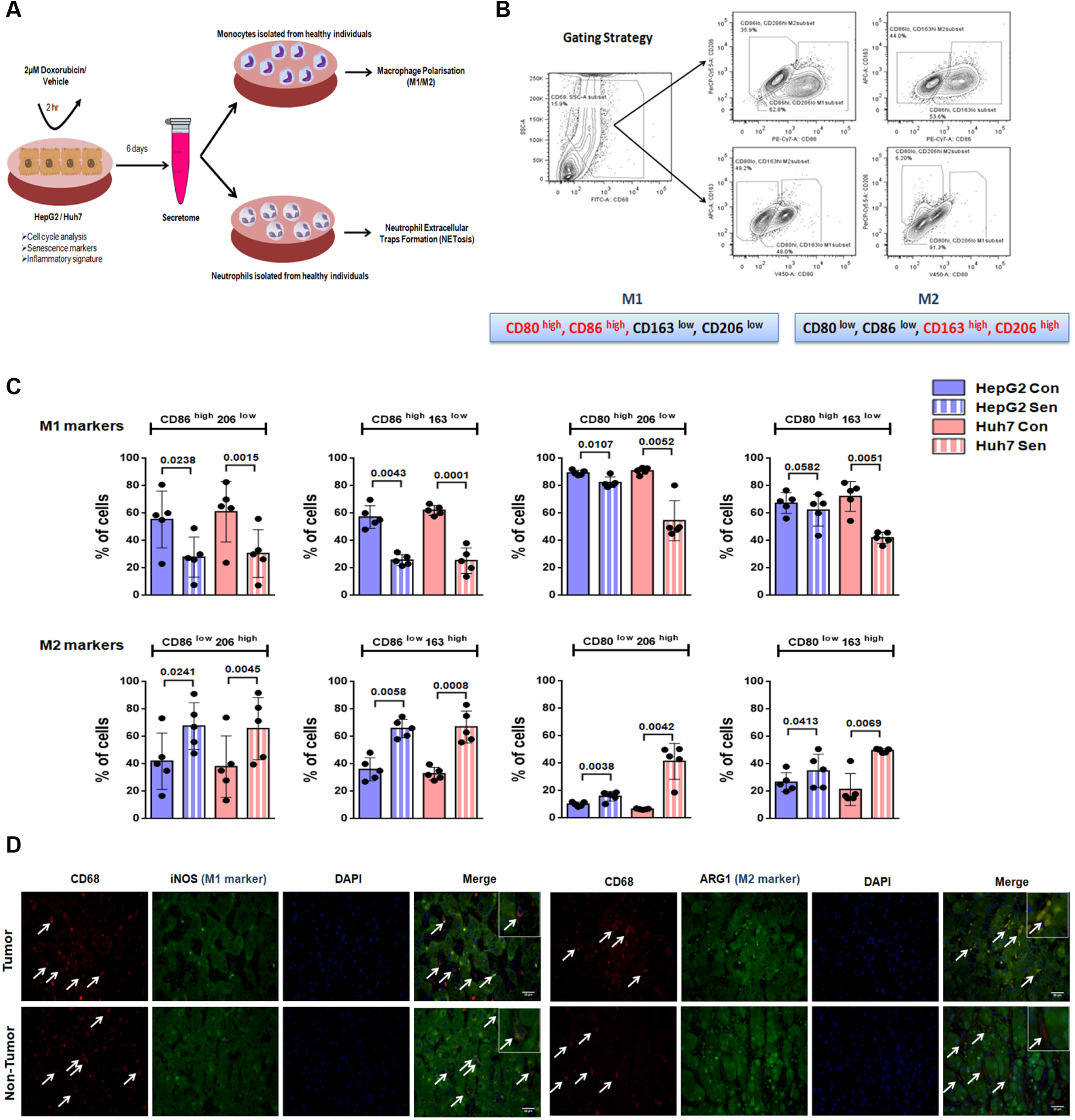
Senescent secretome drive M2 macrophage polarisation. (A) Experimental design for treatment of monocytes and neutrophils isolated from healthy controls with secretome from control and senescent hepatoma cells. (B) Gating strategy for identification of M1 and M2 type of macrophage differentiation. Data was analysed on CD68 gated population. (C) Frequency of M1 (CD86^high^206^low^, CD86^high^163^low^, CD80^high^206^low^, CD80^high^163^low^) and M2 (CD86^low^206^high^, CD86 ^low^163 ^high^, CD80 ^low^206 ^high^, CD80 ^low^163 ^high^) macrophage subtypes in monocytes isolated from healthy individuals and cultured in Control (Con) or senescent (Sen) secretome (N=5). Data is represented as mean ± SD. P values were calculated by paired Student’s t test. (D) Representative immunofluorescence images of CD68 (red), iNOS (green, M1 marker) and arginase1 (green, M2 marker) in tumor and adjacent non-tumor liver tissue. Nucleus was counterstained with DAPI (blue). Merged images show co-localisation of CD68 with iNOS in non-tumor and with arginase1 in tumor region. A zoomed in area is shown in the inset.

### SASP modulates neutrophil extracellular trap formation (NETosis)

Neutrophil extracellular traps (NETs) are formed when neutrophil release chromatin which in turn can modulate tumor growth [25]. Senescent secretome from p53 mutant Huh7 led to an increase in extracellular chromatin and NETs formation; whereas, senescent secretome from HepG2, with a wild type p53, led to a reduction in NETs formation as compared to respective controls (shown in Fig. 4B).

**Fig. 4.**
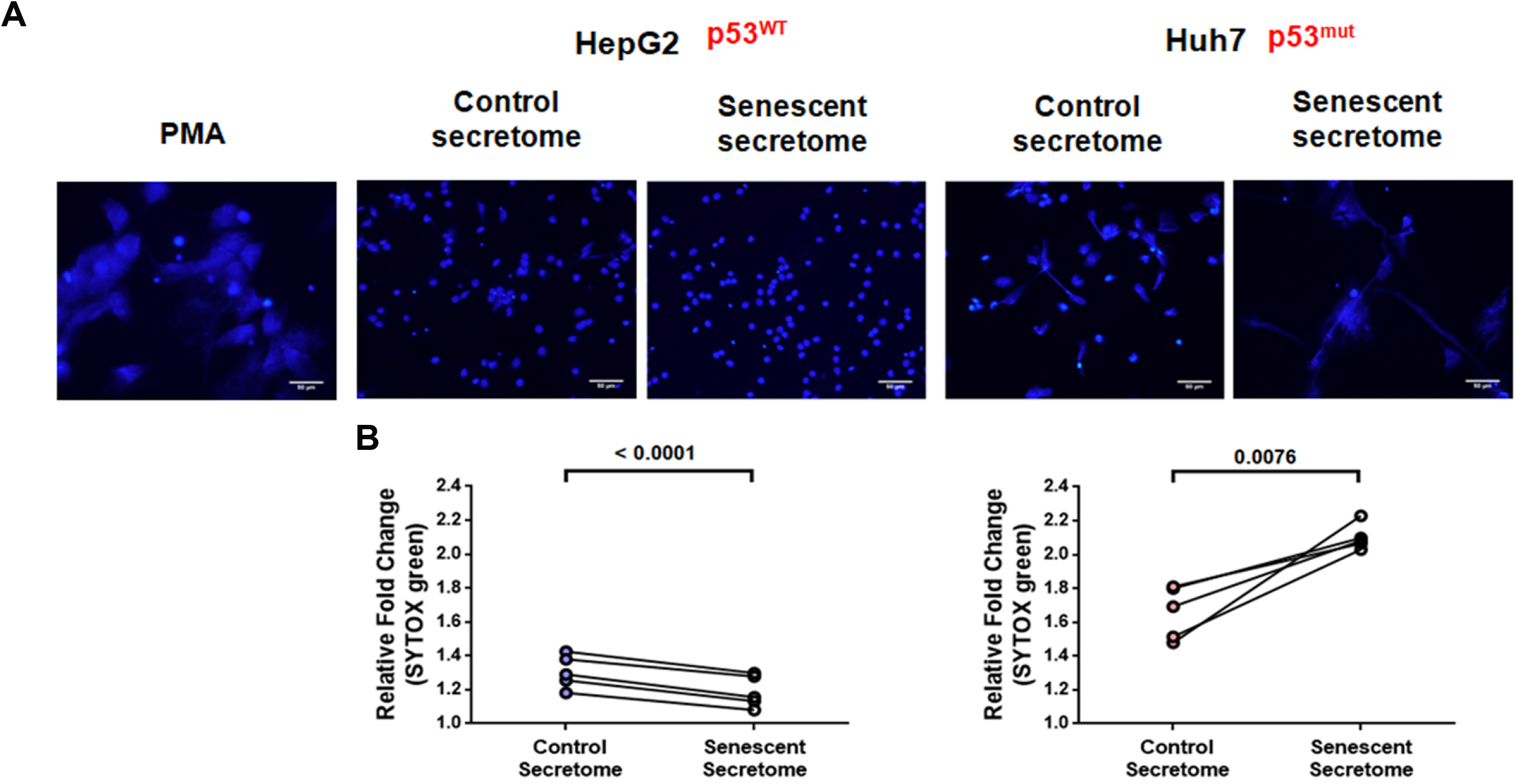
Senescent secretome modulates NETosis. NETosis in neutrophils isolated from healthy controls and treated with either control or senescent secretome of hepatoma cells. (A) Immunofluorescence images of neutrophils (DAPI stained) isolated from healthy individuals and treated with either with PMA (inducer of NETosis) alone or in presence of secretome derived from control or senescent hepatoma cells. (B) Fluorimetry analysis of neutrophils stained with SYTOX Green (dead cells) and Hoechst (live cells). Results were calculated as relative fluorescence units (RFU = SYTOX Green/Hoechst × 100) and data is expressed as relative fold change with respect to untreated controls. Data is represented as mean ± SD of 5 healthy individuals. P values were calculated by paired Student’s t test.

## Discussion

The present work not only corroborated our previous findings on presence of senescent hepatocytes in cirrhosis [9] but also revealed a unique aspect on presence of senescent hepatocytes within the liver tumor microenvironment. Additionally it also highlights the significance of a senescent hepatocyte with an active secretome in a cancerous background in modulating immune cell fate. Senescent cells can chemoattract immune cells for its self clearance which is termed as senescence-surveillance [26]. They can also promote tumorigenesis, (a) by attracting the immature myeloid cells which inhibits the NK cells from tumor cell killing [27] and (b) in obese mice with gut dysbiosis, stellate cell senescence can induce hepatocyte transformation via an active secretory phenotype [28]. While the studies described in literature mostly focused on role of SASP in chemoattraction, there are hardly any studies which have elucidated the influence of SASP in modulating the immune cell fate. Our work demonstrates that senecsnet secretome of HepG2 and Huh7 cells display a pro-inflammatory phenotype, further the. CCR7 and CXCR4, which are predominantly expressed on immune cells, were also increased on the senescent HCC cells implying their proneness towards invasiveness [29–33]. Interestingly, Huh7 with a p53 mutation mounted a stronger inflammatory response than HepG2 with a wild type p53, thereby implicating a role of wild type p53 in modulating tumor inflammation [20].

Macrophages undergo polarisation and the classically activated M1 macrophages are tumoricidal and participate in host defence. Whereas, alternatively activated M2 macrophages produce anti-inflammatory cytokines and are immunosuppressive [34]. Present work highlighted that chemotherapy induced senescence in hepatoma cells was accompanied with a “pro-inflammatory” SASP which in turn drove an “anti-inflammatory” cytokine response through polarisation of macrophages towards an M2 fate. This possibly implicates an unfavourable consequence of senescence towards tumor promotion. Infact, the non-tumorous liver tissue showed presence of predominantly M1 macrophages while the tumor area in the liver showed prevalence of M2 macrophages.

The present work for the first time implicates the role of SASP in the formation of NETs in a p53 dependent manner. In our study, the secretome from Huh7 with a mutant p53 displayed an increase in NETosis when compared with HepG2 (wild type p53). HCC often shows mutations in p53 gene [35]. Often presence of mutant p53 makes the cancer more aggressive. Infact recent studies have indicated a role of NETs in tumor progression and also in initiating a premetastatic niche for cancer cell dissemination [36]. Thus it is possible that SASP induced NETosis may favour tumor cell metastasis in liver cancer and this needs to be explored further.

In conclusion, our study indicates that senescent hepatocyte by eliciting a SASP response can modulate macrophage polarization and NETosis thereby favouring the process of tumorigenesis., Targeting the inflammatory nature of senescent cells, without affecting the growth suppressive features of senescent cells, appears as a promising anti-tumor strategy in future.

## Declarations

## Acknowledgments

BS is a recipient of UGC senior research fellowship. We thank DST-FIST for infrastructural support. We are thankful to Dr. Ritesh Kumar Tiwari (CU-BD, CoE, CRNN) for helping with flow cytometry analysis.

## Availability of data and materials

The datasets generated and/or analyzed during this study are available from the corresponding author on reasonable request.

## Authors Contribution

BS: Performed all experiments, compiled and analysed data, and inputs in manuscript writing, GR: Conceived and designed study, analysed data and wrote manuscript, SA, RN, RS: helped in experimental protocols; AR: Read pathology slides and gave clinical inputs, NT: Helped in FACS and provided research inputs; SKS: Clinical inputs and manuscript editing.

## Ethics approval and consent to participate

The study was approved by the Ethics Committee of Institue of Liver and Biliary Sciences, Delhi, India and written consent of healthy volunteers who consented to give their blood were included in this study.

---

**Fig. S1. Doxorubicin induces growth arrest similar to senescence in hepatoma cells (Huh7 and HepG2).** (A) Bright field microscopy images showing enlarged cells following doxorubicin treatment which stained positive for SA-β-galactosidase assay as indicated by the blue stain. (B) Growth kinetics of hepatoma cells treated with 2μM doxorubicin (Dox) for 2 hr (Day 0), replenished with fresh medium and growth was monitored at specified time intervals (1 to 6 days) by staining with crystal violet dye whose absorbance was checked spectrophotometrically at 570nm. Cells treated with vehicle alone served as control (Con). (C) Cell cycle distribution of control and doxorubicin (Dox) treated cells as analyzed by flow cytometry following EdU incorporation. Following treatment with doxorubicin for 2hr, cells were cultured in fresh medium. On the 6 day cells were pulsed with EdU for 4 hr and then stained with Propidium Iodide (PI) and subjected to flow cytometry for cell cycle analysis. Representative contour plots showing EdU incorporation (y-axis) vs. DNA staining by PI (x-axis). The red box indicate “S-phase cells” which gated positive for EdU and the adjoining bar graph shows the distribution of cells in different phase of cell cycle. The data shown are mean ± SD from three separate experiments. (D) Merged confocal images showing expression of senescence marker LaminB1 (red) in control and doxorubicin treated cells on 6th day. Cells were counterstained with DAPI (blue).

## References

[1] G. Ramakrishna, A. Rastogi, N. Trehanpati, B. Sen, R. Khosla, S.K. Sarin, From cirrhosis to hepatocellular carcinoma: new molecular insights on inflammation and cellular senescence, Liver cancer 2 (2013) 367–383. 10.1159/000343852.

[2] M. Ringelhan, D. Pfister, T. O’Connor, E. Pikarsky, M. Heikenwalder, The immunology of hepatocellular carcinoma, Nature immunology 19 (2018) 222–232. 10.1038/s41590-018-0044-z.

[3] K. Tarao, S. Ohkawa, Y. Miyagi, S. Morinaga, K. Ohshige, N. Yamamoto, M. Ueno, S. Kobayashi, R. Kameda, S. Tamai, Y. Nakamura, K. Miyakawa, Y. Kameda, M. Okudaira, Inflammation in background cirrhosis evokes malignant progression in HCC development from HCV-associated liver cirrhosis, Scandinavian journal of gastroenterology 48 (2013) 729–735. 10.3109/00365521.2013.782064.

[4] A Bishayee, The role of inflammation and liver cancer, Advances in experimental medicine and biology 816 (2014) 401–435. 10.1007/978-3-0348-0837-8_16.

[5] S.U. Wiemann, A. Satyanarayana, M. Tsahuridu, H.L. Tillmann, L. Zender, J. Klempnauer, P. Flemming, S. Franco, M.A. Blasco, M.P. Manns, K.L. Rudolph, Hepatocyte telomere shortening and senescence are general markers of human liver cirrhosis, FASEB journal: official publication of the Federation of American Societies for Experimental Biology 16 (2002) 935–942. 10.1096/fj.01-0977com.

[6] H. Jiang, Z. Ju, K.L. Rudolph, Telomere shortening and ageing, Zeitschrift fur Gerontologie und Geriatrie 40 (2007) 314–324. 10.1007/s00391-007-0480-0.

[7] R.R. Plentz, Y.N. Park, A. Lechel, H. Kim, F. Nellessen, B.H. Langkopf, L. Wilkens, A. Destro, B. Fiamengo, M.P. Manns, M. Roncalli, K.L. Rudolph, Telomere shortening and inactivation of cell cycle checkpoints characterize human hepatocarcinogenesis, Hepatology 45 (2007) 968–976. 10.1002/hep.21552.

[8] A.D. Aravinthan, G.J.M. Alexander, Senescence in chronic liver disease: Is the future in aging?, Journal of hepatology 65 (2016) 825–834. 10.1016/j.jhep.2016.05.030.

[9] B. Sen, A. Rastogi, R. Nath, S.M. Shasthry, V. Pamecha, S. Pandey, K.J. Gupta, S.K. Sarin, N. Trehanpati, G. Ramakrishna, Senescent Hepatocytes in Decompensated Liver Show Reduced UPR(MT) and Its Key Player, CLPP, Attenuates Senescence In Vitro, Cellular and molecular gastroenterology and hepatology 8 (2019) 73–94. 10.1016/j.jcmgh.2019.03.001.

[10] M. Demaria, M.N. O’Leary, J. Chang, L. Shao, S. Liu, F. Alimirah, K. Koenig, C. Le, N. Mitin, A.M. Deal, S. Alston, E.C. Academia, S. Kilmarx, A. Valdovinos, B. Wang, A. de Bruin, B.K. Kennedy, S. Melov, D. Zhou, N.E. Sharpless, H. Muss, J. Campisi, Cellular Senescence Promotes Adverse Effects of Chemotherapy and Cancer Relapse, Cancer discovery 7 (2017) 165–176. 10.1158/2159-8290.CD-16-0241.

[11] K.M. Irvine, R. Skoien, N.J. Bokil, M. Melino, G.P. Thomas, D. Loo, B. Gabrielli, M.M. Hill, M.J. Sweet, A.D. Clouston, E.E. Powell, Senescent human hepatocytes express a unique secretory phenotype and promote macrophage migration, World journal of gastroenterology 20 (2014) 17851–17862. 10.3748/wjg.v20.i47.17851.

[12] A. Lujambio, L. Akkari, J. Simon, D. Grace, D.F. Tschaharganeh, J.E. Bolden, Z. Zhao, V. Thapar, J.A. Joyce, V. Krizhanovsky, S.W. Lowe, Non-cell-autonomous tumor suppression by p53, Cell 153 (2013) 449–460. 10.1016/j.cell.2013.03.020.

[13] D.J. van der Windt, V. Sud, H. Zhang, P.R. Varley, J. Goswami, H.O. Yazdani, S. Tohme, P. Loughran, R.M. O’Doherty, M.I. Minervini, H. Huang, R.L. Simmons, A. Tsung, Neutrophil extracellular traps promote inflammation and development of hepatocellular carcinoma in nonalcoholic steatohepatitis, Hepatology 68 (2018) 1347–1360. 10.1002/hep.29914.

[14] J. Claria, R.E. Stauber, M.J. Coenraad, R. Moreau, R. Jalan, M. Pavesi, A. Amoros, E. Titos, J. Alcaraz-Quiles, K. Oettl, M. Morales-Ruiz, P. Angeli, M. Domenicali, C. Alessandria, A. Gerbes, J. Wendon, F. Nevens, J. Trebicka, W. Laleman, F. Saliba, T.M. Welzel, A. Albillos, T. Gustot, D. Benten, F. Durand, P. Gines, M. Bernardi, V. Arroyo, C.S.I.o.t.E.-C. Consortium, F. the European Foundation for the Study of Chronic Liver, Systemic inflammation in decompensated cirrhosis: Characterization and role in acute-on-chronic liver failure, Hepatology 64 (2016) 1249–1264. 10.1002/hep.28740.

[15] A Krtolica, S. Parrinello, S. Lockett, P.Y. Desprez, J. Campisi, Senescent fibroblasts promote epithelial cell growth and tumorigenesis: a link between cancer and aging, Proceedings of the National Academy of Sciences of the United States of America 98 (2001) 12072–12077. 10.1073/pnas.211053698.

[16] T. Kuilman, C. Michaloglou, L.C. Vredeveld, S. Douma, R. van Doorn, C.J. Desmet, L.A. Aarden, W.J. Mooi, D.S. Peeper, Oncogene-induced senescence relayed by an interleukin-dependent inflammatory network, Cell 133 (2008) 1019–1031. 10.1016/j.cell.2008.03.039.

[17] T. Kuilman, C. Michaloglou, W.J. Mooi, D.S. Peeper, The essence of senescence, Genes & development 24 (2010) 2463–2479. 10.1101/gad.1971610.

[18] G.P. Dimri, X. Lee, G. Basile, M. Acosta, G. Scott, C. Roskelley, E.E. Medrano, M. Linskens, I. Rubelj, O. Pereira-Smith, et al., A biomarker that identifies senescent human cells in culture and in aging skin in vivo, Proceedings of the National Academy of Sciences of the United States of America 92 (1995) 9363–9367.

[19] C. Carmona-Rivera, M.J. Kaplan, Induction and Quantification of NETosis, Current Protocols in Immunology 115 (2016) 14.41.11–14.41.14. doi:10.1002/cpim.16.

[20] C.D. Wiley, N. Schaum, F. Alimirah, J.A. Lopez-Dominguez, A.V. Orjalo, G. Scott, P.Y. Desprez, C. Benz, A.R. Davalos, J. Campisi, Small-molecule MDM2 antagonists attenuate the senescence-associated secretory phenotype, Scientific reports 8 (2018) 2410. 10.1038/s41598-018-20000-4.

[21] R.S. Bystry, V. Aluvihare, K.A. Welch, M. Kallikourdis, A.G. Betz, B cells and professional APCs recruit regulatory T cells via CCL4, Nature immunology 2 (2001) 1126–1132. 10.1038/ni735.

[22] K. Beider, M. Abraham, M. Begin, H. Wald, I.D. Weiss, O. Wald, E. Pikarsky, R. Abramovitch, E. Zeira, E. Galun, A. Nagler, A. Peled, Interaction between CXCR4 and CCL20 pathways regulates tumor growth, PloS one 4 (2009) e5125. 10.1371/journal.pone.0005125.

[23] I.S. Afonina, C. Muller, S.J. Martin, R. Beyaert, Proteolytic Processing of Interleukin-1 Family Cytokines: Variations on a Common Theme, Immunity 42 (2015) 991–1004. 10.1016/j.immuni.2015.06.003.

[24] O.W. Yeung, C.M. Lo, C.C. Ling, X. Qi, W. Geng, C.X. Li, K.T. Ng, S.J. Forbes, X.Y. Guan, R.T. Poon, S.T. Fan, K. Man, Alternatively activated (M2) macrophages promote tumour growth and invasiveness in hepatocellular carcinoma, Journal of hepatology 62 (2015) 607–616. 10.1016/j.jhep.2014.10.029.

[25] M. Demers, S.L. Wong, K. Martinod, M. Gallant, J.E. Cabral, Y. Wang, D.D. Wagner, Priming of neutrophils toward NETosis promotes tumor growth, Oncoimmunology 5 (2016) e1134073. 10.1080/2162402X.2015.1134073.

[26] T.W. Kang, T. Yevsa, N. Woller, L. Hoenicke, T. Wuestefeld, D. Dauch, A. Hohmeyer, M. Gereke, R. Rudalska, A. Potapova, M. Iken, M. Vucur, S. Weiss, M. Heikenwalder, S. Khan, J. Gil, D. Bruder, M. Manns, P. Schirmacher, F. Tacke, M. Ott, T. Luedde, T. Longerich, S. Kubicka, L. Zender, Senescence surveillance of pre-malignant hepatocytes limits liver cancer development, Nature 479 (2011) 547–551. 10.1038/nature10599.

[27] T. Eggert, K. Wolter, J. Ji, C. Ma, T. Yevsa, S. Klotz, J. Medina-Echeverz, T. Longerich, M. Forgues, F. Reisinger, M. Heikenwalder, X.W. Wang, L. Zender, T.F. Greten, Distinct Functions of Senescence-Associated Immune Responses in Liver Tumor Surveillance and Tumor Progression, Cancer cell 30 (2016) 533–547. 10.1016/j.ccell.2016.09.003.

[28] S. Yoshimoto, T.M. Loo, K. Atarashi, H. Kanda, S. Sato, S. Oyadomari, Y. Iwakura, K. Oshima, H. Morita, M. Hattori, K. Honda, Y. Ishikawa, E. Hara, N. Ohtani, Obesity-induced gut microbial metabolite promotes liver cancer through senescence secretome, Nature 499 (2013) 97–101. 10.1038/nature12347.

[29] H. Ma, L. Gao, S. Li, J. Qin, L. Chen, X. Liu, P. Xu, F. Wang, H. Xiao, S. Zhou, Q. Gao, B. Liu, Y. Sun, C. Liang, CCR7 enhances TGF-beta1-induced epithelial-mesenchymal transition and is associated with lymph node metastasis and poor overall survival in gastric cancer, Oncotarget 6 (2015) 24348–24360. 10.18632/oncotarget.4484.

[30] L. Yang, Y. Chang, P. Cao, CCR7 preservation via histone deacetylase inhibition promotes epithelial-mesenchymal transition of hepatocellular carcinoma cells, Experimental cell research 371 (2018) 231–237. 10.1016/j.yexcr.2018.08.015.

[31] X. Wang, W. Zhang, Y. Ding, X. Guo, Y. Yuan, D. Li, CRISPR/Cas9-mediated genome engineering of CXCR4 decreases the malignancy of hepatocellular carcinoma cells in vitro and in vivo, Oncology reports 37 (2017) 3565–3571. 10.3892/or.2017.5601.

[32] P.T. Gao, G.Y. Ding, X. Yang, R.Z. Dong, B. Hu, X.D. Zhu, J.B. Cai, Y. Ji, G.M. Shi, Y.H. Shen, J. Zhou, J. Fan, H.C. Sun, C. Huang, Invasive potential of hepatocellular carcinoma is enhanced by loss of selenium-binding protein 1 and subsequent upregulation of CXCR4, American journal of cancer research 8 (2018) 1040–1049.

[33] S. Chatterjee, B. Behnam Azad, S. Nimmagadda, The intricate role of CXCR4 in cancer, Advances in cancer research 124 (2014) 31–82. 10.1016/B978-0-12-411638-2.00002-1.

[34] S. Aras, M.R. Zaidi, TAMeless traitors: macrophages in cancer progression and metastasis, British journal of cancer 117 (2017) 1583–1591. 10.1038/bjc.2017.356.

[35] P.A. Muller, K.H. Vousden, Mutant p53 in cancer: new functions and therapeutic opportunities, Cancer cell 25 (2014) 304–317. 10.1016/j.ccr.2014.01.021.

[36] W. Lee, S.Y. Ko, M.S. Mohamed, H.A. Kenny, E. Lengyel, H. Naora, Neutrophils facilitate ovarian cancer premetastatic niche formation in the omentum, The Journal of experimental medicine 216 (2019) 176–194. 10.1084/jem.20181170.

