## Supplementary figures and images for "Secretome of senescent hepatoma cells modulate macrophage polarization and neutrophil extracellular traps formation"

### Supplemental file

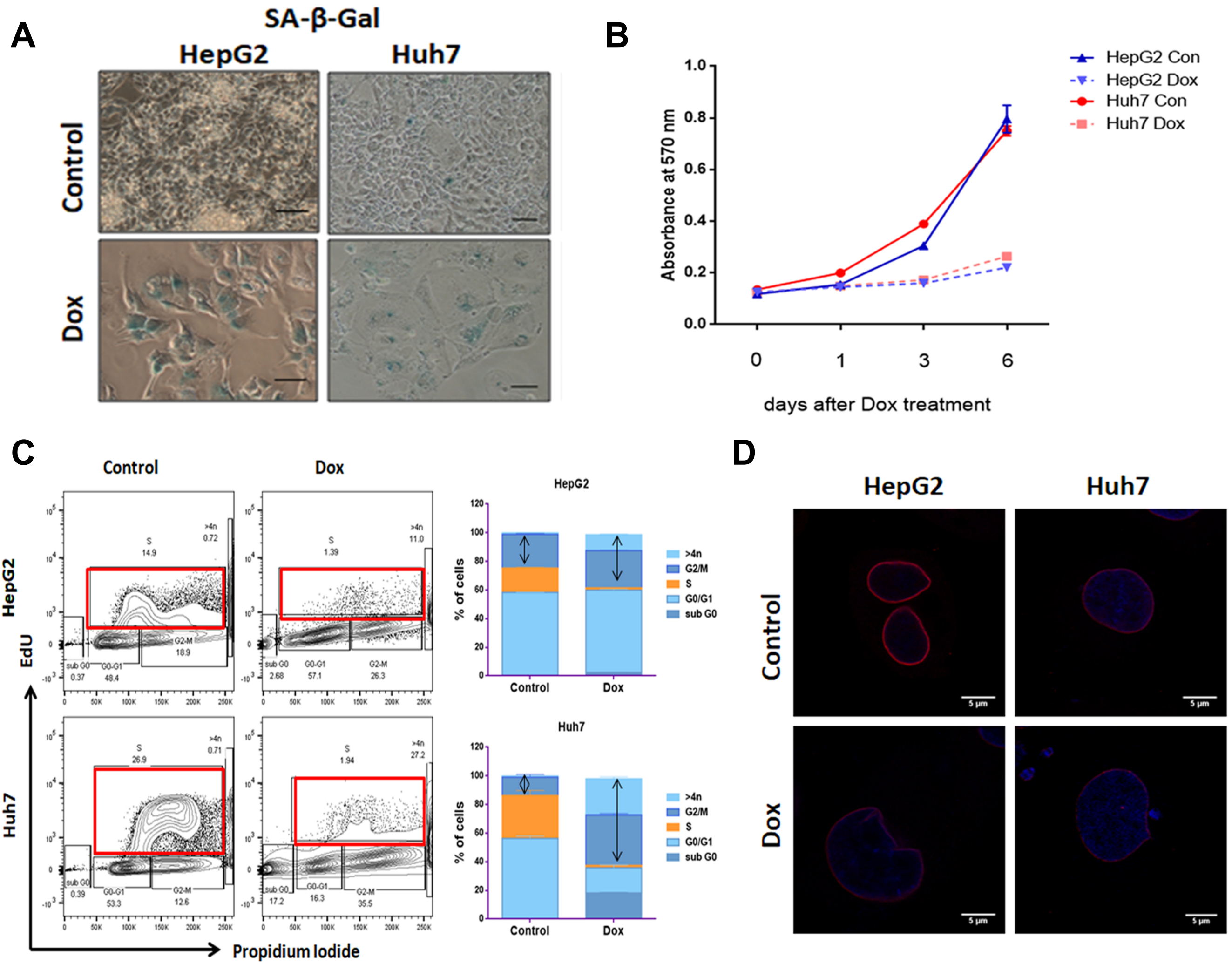
